# Human tauopathies are not associated with an activated unfolded protein response

**DOI:** 10.1101/2019.12.20.884437

**Authors:** A. P. Pitera, I. J. Hartnell, D. Boche, V. O’Connor, K. Deinhardt

## Abstract

Tauopathies are the neurodegenerative diseases associated with the accumulation of misfolded tau protein. Despite many years of investigation, the mechanisms underpinning tau dependent proteinopathy remains to be elucidated. A protein quality control pathway within the endoplasmic reticulum (ER), called unfolded protein response (UPR), has been suggested as a possible response implicated in the misfolded tau-mediated neurodegeneration. However, the question arose: how does the cytosolic protein tau that does not enter the ER induce a response stemming from this compartment? In this study we investigated three different human tauopathies to establish whether these diseases are associated with the activation of UPR. We probed for the modulation of several reliable UPR markers in mRNA and proteins extracted from 20 brain samples from Alzheimer’s disease (AD) patients, 11 from Pick’s disease (PiD) and 10 from Progressive Supranuclear Palsy (PSP) patients coupled to equal numbers of age-matched non-demented controls. This showed that different markers of UPR are not changed in any of the human tauopathies investigated. Interestingly, UPR signatures were often observed in non-demented controls. These data from human tissue further support the emerging evidence that the accumulation of misfolded cytosolic tau does not drive a diseased associated activation of UPR.

## Introduction

The microtubule associated protein tau is misfolded and accumulated in several conditions, including the most common cause of dementia, Alzheimer’s disease (AD) [1]. In AD, the accumulation of tau lesions forms neurofibrillary tangles (NFTs) and is accompanied by the deposition of β-amyloid (Aβ) plaques [2]. The presence of Aβ plaques alone is not sufficient to cause neurodegeneration, and Aβ deposition has been reported in aged brains without the presence of clinical symptoms of dementia [3,4]. In contrast, tau pathology in AD is well correlated with the cognitive and clinical symptoms. Further, deposition of tau can be observed in other neurodegenerative diseases without the Aβ plaques influence [5]. These diseases can be classified as primary tauopathies due to the fact that tau is a primary hallmark of the observed pathology [2]. Different tauopathies are distinguished by the relative contribution or prevalence of the distinct tau isoforms (3R or 4R) to the misfolded tau deposits. Furthermore, these distinct molecular entities are associated with differential anatomical distribution of the disease affected cell types [6]. Pick’s disease (PiD) is a 3R tauopathy in which tau accumulates forming large spherical Pick bodies. In contrast Progressive Supranuclear Palsy (PSP) is an example of a 4R tauopathy with the spherical ‘globose’ NFTs being found in degenerating neuronal structures [7]. Importantly, high resolution structural information of the purified aggregates from distinct diseases indicates the presence of disease selective misfolds [8,9].

There are currently no disease-modifying treatments against neurodegenerative diseases. However, the fact that neurodegeneration is associated with disturbed protein homeostasis is well established [10,11]. For this reason, the unfolded protein response (UPR), a protein quality control pathway within the endoplasmic reticulum (ER), is frequently suggested to be a mechanism implicated in neurodegenerative processes. The UPR is induced as a response to protein misfolding occurring within the lumen of the ER. Three branches of an activated UPR can be distinguished and characterised by different signalling components: IRE1 (inositol requiring enzyme 1), PERK (protein kinase R-like endoplasmic reticulum kinase) and ATF6 (activating transcription factor 6) [12,13]. Activation of UPR increases the protein-folding capacity and re-establishes protein homeostasis. However prolonged UPR activation leads to cellular death [12]. UPR induction has been described for different neurodegenerative diseases, including tauopathies such as AD and PiD [14,15]. Therefore, the idea of targeting UPR as a possible treatment for tau-related neurodegenerative diseases has emerged [16,17]. However, this concept has been called into question due to the fact that tau is a cytosolic protein that does not form aggregates in the ER compartment and thus it is unclear how tau misfolding would induce a UPR activation [11].

UPR induction has been investigated in transgenic mouse models of tauopathy with opposing results being reported [16–18]. We have previously used *in vivo* and *in vitro* pre-clinical models of tauopathy and found no indication of UPR activation in response to misfolded tau [19]. Since these models recapitulate only select aspects of the human pathology, we investigated whether human tauopathies evidence an activation of the UPR. We performed qPCR and Western blot experiments using the reagents that readily detect the UPR in the perturbed human cell lines. We measured the mRNA and protein level of several markers involved in the UPR branches in human from three different diseases related to tau pathology and found that tau pathology is not associated with UPR activation. This study utilizing human samples is consistent with a growing number of studies that do not report UPR in experimental models of chronic neurodegeneration [18,19].

## Materials and methods

### Materials

The primary antibodies used in this study were: anti-BiP (Cell Signaling, #3177, 1:500 dilution), anti-CHOP (Cell Signaling, #2896, 1:1000 dilution), anti-p-eIF2α (Cell Signaling, #3398, 1:1000 dilution), anti-tau (DAKO, #A0024, 1:1000 dilution), anti-GAPDH (Abcam, #ab8245, 1:5000 dilution) and PHF-1 (gift from Prof. Peter Davies, 1:5000 dilution, [20]). Secondary antibodies were conjugated to IRDyes (LI-COR Biosciences) and used at 1:10000 dilution. The oligonucleotide primers used in the qPCR and PCR reactions were obtained from Eurofins Genomics. The sequences of the primers for each gene are shown in Table 1.

**Table 1.**
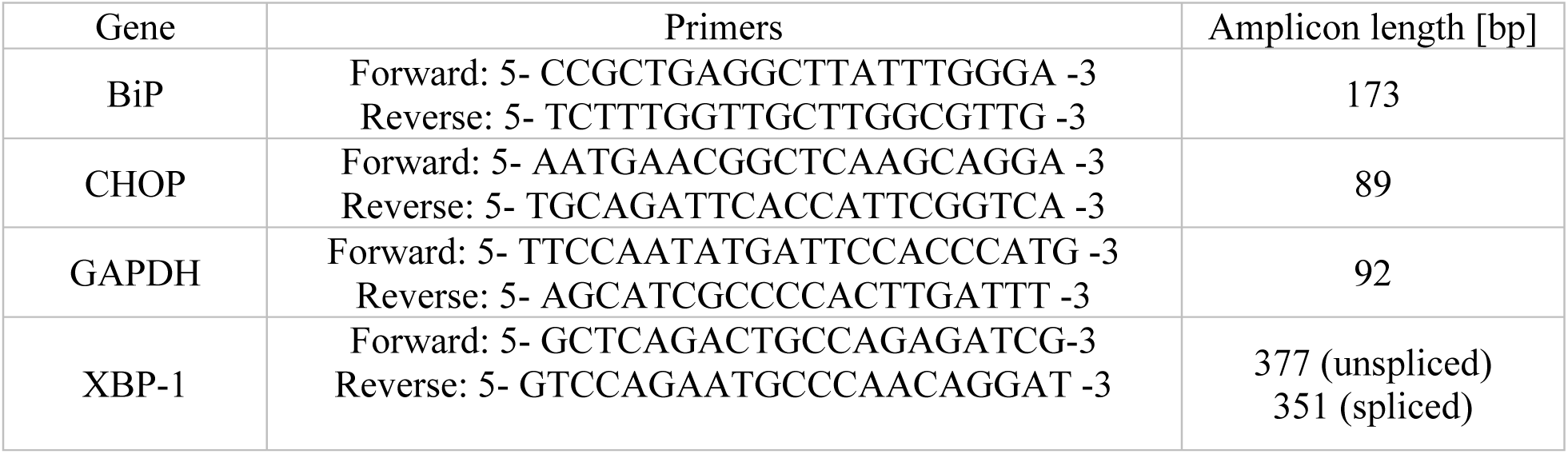
Sequences of the primers used for the PCR and qPCR experiments. bp-base pair.

### Cases

AD brain tissue was sourced from the South West Dementia Brain Bank comprising 20 AD cases and 20 controls (nAD). The inferior parietal lobule, as an area of the cerebral cortex typically affected by AD pathology [21], was investigated for all cases (Table 2). Eleven PiD and 11 controls (nPiD) as well as 10 PSP and 10 controls (nPSP) were provided by the London Neurodegenerative Diseases Brain Bank, the South West Dementia Brain Bank and the Oxford Brain Bank with the frontal cortex investigated for the FTD and control cases (Table 3 for nPiD and PiD cases and Table 4 for nPSP and PSP cases). Cases with any other significant brain pathologies such as stroke, tumour, or traumatic brain injury were excluded from the study. Controls with no history of neurological or psychiatric disease or symptoms of cognitive impairment were matched with age, gender and *post-mortem* delay as closely as possible. To minimize the time in formalin, which has an effect on the quality of the immunostaining, the selection was performed on the availability of formalin fixed paraffin embedded tissue, and thus on blocks processed at the time of the original *post-mortem* examination. Fresh frozen tissue with a pH > 5.5 to ensure RNA integrity [22,23] was used for detection of mRNA by qPCR and proteins by Western blot.

**Table 2.** Demographic, clinical and post-mortem characteristics of nAD and AD groups. N/A-non-applicable.

| COHORTS | nAD<br>N=20 | AD<br>N=20 |
| --- | --- | --- |
| **GENDER** | 13F:7M | 11F:9M |
| **AGE OF DEATH**<br>**(YEARS, MEAN ± SD)** | 85±1.3 | 80±1.3 |
| **AGE OF AD ONSET**<br>**(YEARS, MEAN ± SD)** | N/A | 70±1.4 |
| **DURATION OF AD**<br>**(YEARS, MEAN ± SD)** | N/A | 10±1.0 |
| **BRAAK STAGE** | 0-II: 19<br>III-IV: 1<br>V-VI: 0 | 0-II: 0<br>III-IV: 2<br>V-VI: 18 |
| **APOE GENOTYPE** | ε4/- : 2<br>ε4/ε4: 1 | ε4/- : 3<br>ε4/ε4: 3 |
| **POST-MORTEM DELAY**<br>**(HOURS, MEAN ± SD)** | 43±6 | 43±6 |

**Table 3.**
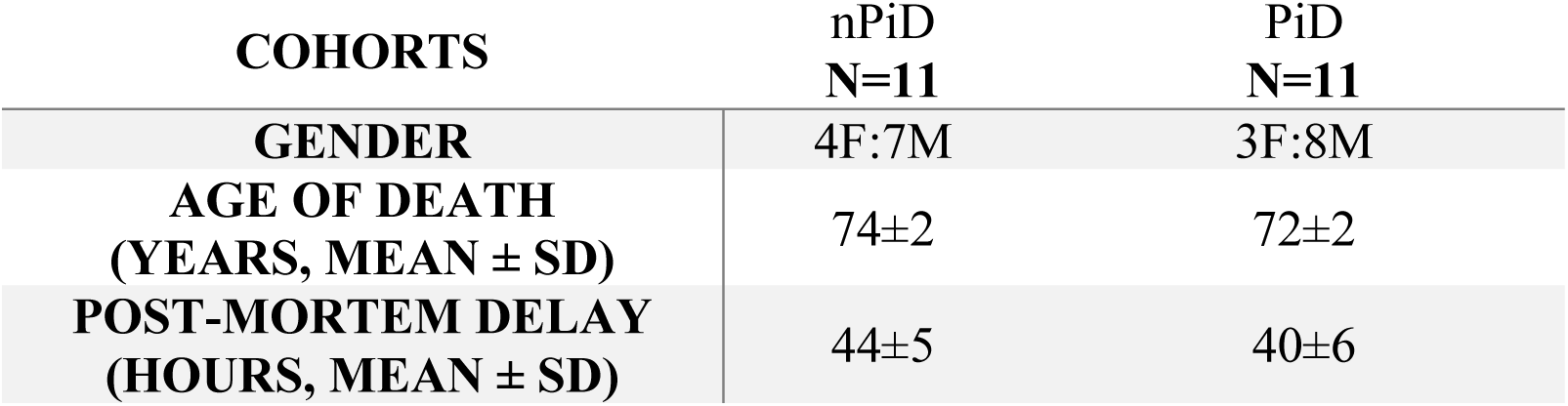
Demographic, clinical and post-mortem characteristics of nPiD and PiD groups.

**Table 4.** Demographic, clinical and post-mortem characteristics of nPSP and PSP groups.

| **COHORTS** | **nPSP**<br>**N=10** | **PSP**<br>**N=10** |
| --- | --- | --- |
| **GENDER** | 6F:4M | 5F:5M |
| **AGE OF DEATH**<br>**(YEARS, MEAN ± SD)** | 77±2 | 75±2 |
| **POST-MORTEM DELAY**<br>**(HOURS, MEAN ± SD)** | 39±6 | 37±5 |

### Ethics

Ethical approval was provided by the South West Dementia Brain Bank (REC approval 08/H0106/28+5, the London Neurodegenerative Diseases Brain Bank (REC approval 08/H0704/128), and the Oxford Brain bank (REC approval 51/SC/0639).

### Immunohistochemistry

Immunohistochemistry against phosphorylated (p)-tau protein was performed on 6 μm paraffin-embedded sections using the clone AT8 (#MN1020, ThermoScientific). Biotinylated secondary antibody goat anti-mouse was from Vector Laboratories (Peterborough, UK). Presence of p-tau was visualized using the avidin–biotin–peroxidase complex method (Vectastain Elite, Vector Laboratories) with 3,3′-diaminobenzidine as chromogen and 0.05% hydrogen peroxide as substrate (Vector Laboratories). All sections were counterstained with haematoxylin, then dehydrated and mounted in Pertex (Histolab Products AB). Staining was performed in several batches with each batch containing disease and control cases to ensure comparability of immunolabelling. All experiments included a negative control slide incubated in buffer with no primary antibody.

### Quantification

Quantification was blinded to the case designation. For each case, 30 images of grey matter were acquired by the Olympus dotSlide virtual microscopy system under a x20 objective. The images were obtained in a zigzag sequence to ensure sampling of all six cortical layers as previously published [24,25]. Quantitative image analysis was carried out using ImageJ (version 1.52p, Wayne Rasband, NIH, USA). A specific threshold was determined to quantify the area fraction of each image labelled by AT8 and expressed as protein load (%), and the mean value was calculated for each case.

### Tissue homogenisation

The brain samples were homogenised in 5 volumes (w/v) of sterile PBS containing cOmplete Mini EDTA-free Protease Inhibitor Cocktail tablets (Roche), sodium fluoride (10 mM, Fisher Scientific) and sodium orthovanadate (2 mM, Fisher Scientific). The samples were homogenised using Kontes pellet pestle motor and plastic pellet pestles. The homogenised samples in PBS were then used for RNA or protein extraction [26].

### qPCR

RNA was isolated from 100 µl of homogenate using the Trizol method. The RNA was purified using RNeasy kit (Qiagen). RNA amount and quality was analysed using NanoDrop spectrophotometer (Thermo Fisher Scientific). The RNA exhibited an absorbance maximum at 260 nm (A260) and the ratios of A260/A280 and 260/230 were ∼2. The extracted RNA was treated with Precision DNase (Primer Design) according to manufacturer’s instructions. 200 ng of RNA was reverse transcribed using Precision nanoScript 2 reverse transcription kit (Primer Design) according to manufacturer’s instructions.

qPCR was performed using Precision PLUS Mastermix (Primer Design) according to the manufacturer’s instructions. The mix containing 1 µl of cDNA (obtained as described above), solution of forward and reverse primer (0.2 µM), Precision PLUS Mastermix StepOnePlus and nuclease-free water was incubated in the Real-Time PCR instrument (Applied Biosystems) using the following conditions: hot start 95**°**C for 2 min and 40 cycles of denaturation at 95**°**C for 10 s and data collection for 1 min.

All reactions were performed with two technical replicates. Standard curves for each set of primers were generated using cDNA from HEK293 cells. A cDNA serial dilution curve was constructed from 5-fold serial dilutions as follows: neat cDNA, 1:5, 1:25, 1:125 and 1:625 dilutions. The expression level was normalized to the level of the reference gene, GAPDH. To ensure the amplification of desired amplicon without unspecific products, a melting curve was performed for each qPCR reaction and qPCR products were run on 2% agarose gel. qPCR reaction with no template was run alongside the reactions with the cDNA samples to exclude the possibility of contamination.

### XBP-1 splicing

RedTaq ReadyMix PCR Reaction Mix (Sigma Aldrich), 1 µl of cDNA (obtained as described above) and primers (0.2 µM) detecting both, spliced and unspliced, forms of XBP-1 were used for end point PCR. The samples were kept on ice before being heated at 94**°**C for 2 min. Subsequently, the samples were subjected to 40 cycles of denaturation at 94**°**C for 40 s, annealing at 60**°**C for 30 s, extension at 72**°**C for 1 min using GeneAmp PCR System 9700 (Applied Biosystems). A final extension was performed at 72**°**C for 10 min. The amplicons of this high cycle number PCR reaction were separated on 2.5% agarose gel. The no template control that did not contain cDNA was run alongside the samples and failed to show XBP-1 amplification. It shows the specificity of the amplification.

### Western blotting

Proteins from PBS-homogenised samples were extracted with the equal volume of 2x extraction buffer containing HEPES-NaOH (40 mM, pH 7.4, Fisher Scientific), NaCl (250 mM, Sigma Aldrich), SDS (4%), cOmplete Mini EDTA-free Protease Inhibitor Cocktail tablets (Roche), sodium fluoride (10 mM) and sodium orthovanadate (2 mM). After extraction, protein concentration was measured using Bio-Rad DC protein assay kit according to manufacturer’s instructions.

The samples were mixed to give a final concentration of 2 µg/µl with 5x sample buffer containing tris-HCl (312.5 mM, pH 6.8, SDS (10%), glycerol (50%), dithiothreitol (25 mM) and bromophenol blue dye (0.005%).

These samples were boiled for 10 min at 95**°**C, briefly spun before loading to 12% acrylamide gel. To minimise the risk of inconsistent analysis of tauopathy cases and non-demented controls due to the differences in gels/gel transfer, the patient samples were always run on the same gel as the relative age-matched controls. After loading, the samples were separated by SDS-PAGE and transferred to nitrocellulose membrane. The membranes were blocked in the blocking solution (2.5% BSA, TBS with 0.1% Tween) for 1h and then incubated at 4**°**C overnight with the primary antibody diluted in the blocking solution. The membrane was then incubated with the secondary antibody at room temperature for 1h. Immunoreactivity was revealed using an Odyssey Infrared Imaging System (LI-COR Biosciences). The Image Studio Scanner software was used to capture the image and Image Studio Lite software was used to quantify the intensities of the bands. Only the antibodies that were capable of detecting increased immunoreactivity of positive control were used in this study.

### Statistical analysis

The normality of distribution of the data was determined using Schapiro-Wilk test. Potential differences in the expression of the UPR markers between disease and control cases were compared using either T-test for parametric data or Mann-Whitney U-test for non-parametric data. One-sided tests were used when comparing the level of p-tau between disease and control cases based on the prediction that p-tau is increased in tauopathy cases. For the other UPR markers, two-sided tests were assumed. Correlations between the different markers were analysed by either Pearson’s (parametric) or Spearman’s (non-parametric) test. Fisher’s exact test was used for comparison between disease and control cases to assess the presence of XBP-1 splicing. All analyses were performed with SPSS software (version 25, IBM). P values less than 0.05 for intergroup comparisons and 0.01 for correlations were considered statistically significant. Graphs were prepared with GraphPad Prism software (version 8, La Jolla, CA).

### Data availability

All data generated or analysed during this study are included in this published article and its supplementary information files.

## Results

### Tau phosphorylation in neurodegeneration

Tauopathies are characterised by the increased phosphorylation and accumulation of hyperphosphorylated tau. We used immunohistochemistry to investigate tauopathy and confirmed the presence of tau pathology in the AD, PiD and PSP cases, using the commonly utilised AT8 antibody (recognising pSer202/pThr205) (Figure 1A). In AD, dystrophic neurites, neuropil threads and neurofibrillary tangles (NFTs) were observed. For PiD p-tau was localised in neurons, labelling Pick bodies, and neuropil. In the PSP cases the pathology clearly presented NFTs. Representative stains that highlight these features are shown. In parallel, we homogenised and extracted mRNA and proteins from the brains of cases listed (Methods section). To benchmark the cases investigated, we performed Western blot and used PHF1 antibody directed against phospho-epitopes Ser396/Ser404. The AD, PiD and PSP cases showed increased phosphorylation when compared to the control cases as follows: AD: p<0.0001, PiD: p=0.0006, PSP: p=0.0045 when normalised to GAPDH and AD: p<0.0001, PiD: p=0.0006, PSP: p=0.0010 when normalised to the total tau level (Figure 1B). The increase in tau phosphorylation in AD cases was accompanied by the increase in total tau when compared to age-matched non-demented controls (p=0.0005). In contrast, PiD cases were not characterised by increased total tau level (p=0.2055); whereas an 80-fold increase of p-tau level was detected when compared to controls. Similarly, in PSP cases, total tau level was not increased (p=0.5718) compared to control cases, despite the significantly increased p-tau.

**Figure 1.**
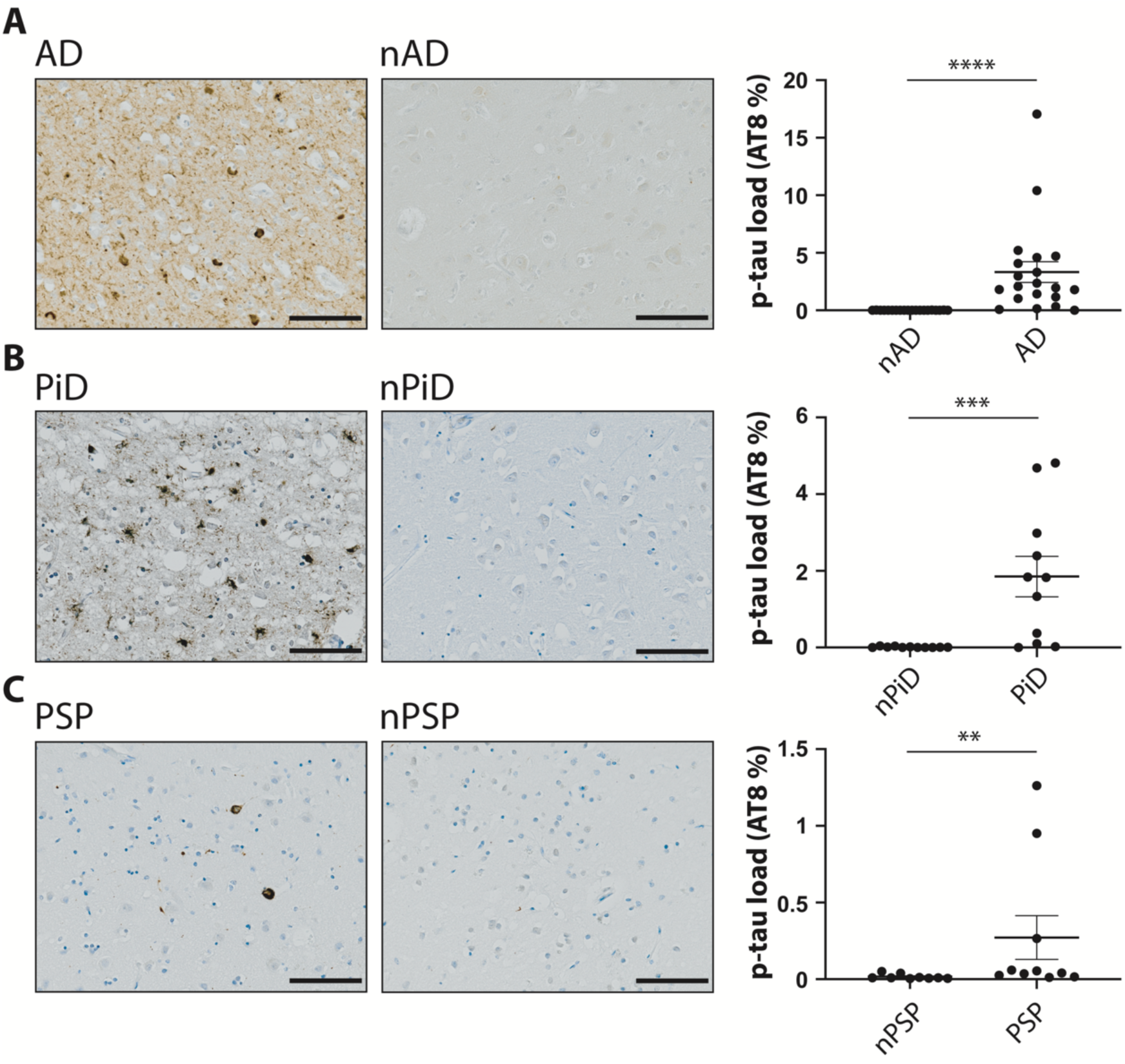
Representative images of p-tau staining with AT8 antibody in brain samples from distinct tauopathy cohorts: AD and age-matched controls nAD (A), PiD and age-matched nPiD (B), PSP and age-matched nPSP (C). The quantification of AT8 staining was analysed using Mann-Whitney test. Error bars are SEM. ****p≤0.0001; ***p≤0.001; **p≤0.01. Scale bar indicates 100 μm.

Interestingly, the phosphorylation level in the cases within the diseased groups (AD, Pi and PSP) was heterogenous showing up to an 80-fold variation in p-tau between the highest and lowest levels of PHF-1 immunoreactivity detected by western blotting. In contrast, the relative levels of total tau showed a much more constrained variability of immunoreactivity. Finally, this side by side comparison of distinct disease harbouring homogenates allowed us to score the relative levels of p-tau. Although increased p-tau was detected for all the diseases, the degree of tau phosphorylation differed amongst the tauopathies, with PSP cases having the lowest fold increase in comparison to non-demented cohort. The data confirm the increased phosphorylation of tau in the brains of patients affected with dementia. What is more, the results also highlight the variability of p-tau level that can be observed within the diseased groups.

### XBP-1 splicing in tauopathy

One of the commonly used and sensitive markers of UPR activation is splicing of XBP-1. This generates an active transcription factor that leads to the induced expression of the genes involved in ER homeostasis [27]. We have previously shown that upon chemical perturbation rodent neurons in cultures undergo a robust UPR, readily detected by primers designed to pick up XBP-1 splicing [19]. We designed human primers that span the spliced intron and enable detection of both forms to determine whether the splicing event occurs in a disease. These primers were first verified on the mRNA extracted from tunicamycin-treated HEK293 cells. This identified that they selectively detected the spliced and unspliced forms of XBP-1 in human cells (Supplement S1A).

The primers amplified the products of expected size in cDNA from human brain samples. We then used the distribution of the spliced and unspliced amplicons to probe if XBP-1 splicing was selectively occurring in the investigated tauopathies when compared to the respective controls (Figure 2). We observed that splicing occurs independently of the presence of the disease diagnosis. In AD cases, 40% of the samples were found to be positive for the splicing compared to 75% in controls. There was no significant difference between both groups (p= 0.1560). In PiD, 55% of the cases presented the XBP-1 splicing but this was not different from the control cases (p=0.9999). Similarly, in PSP, 40% of the cases were positive for XBP-1 splicing with no significant difference when compared to respective control cases (p=0.6372). For each group, some cases had undetectable level of XBP-1 despite readily amplifying the housekeeping gene which suggests that the lack of XBP-1 message was not due to the cDNA quality. Further, there was no association between the XBP-1 splicing and the level of p-tau (data not shown). Altogether, the data suggest that XBP-1 splicing occurs independently from the presence of tauopathy and that it is a widely observed event in the ageing brain based on its presence in the non-demented control samples (Supplement S2).

**Figure 2.**
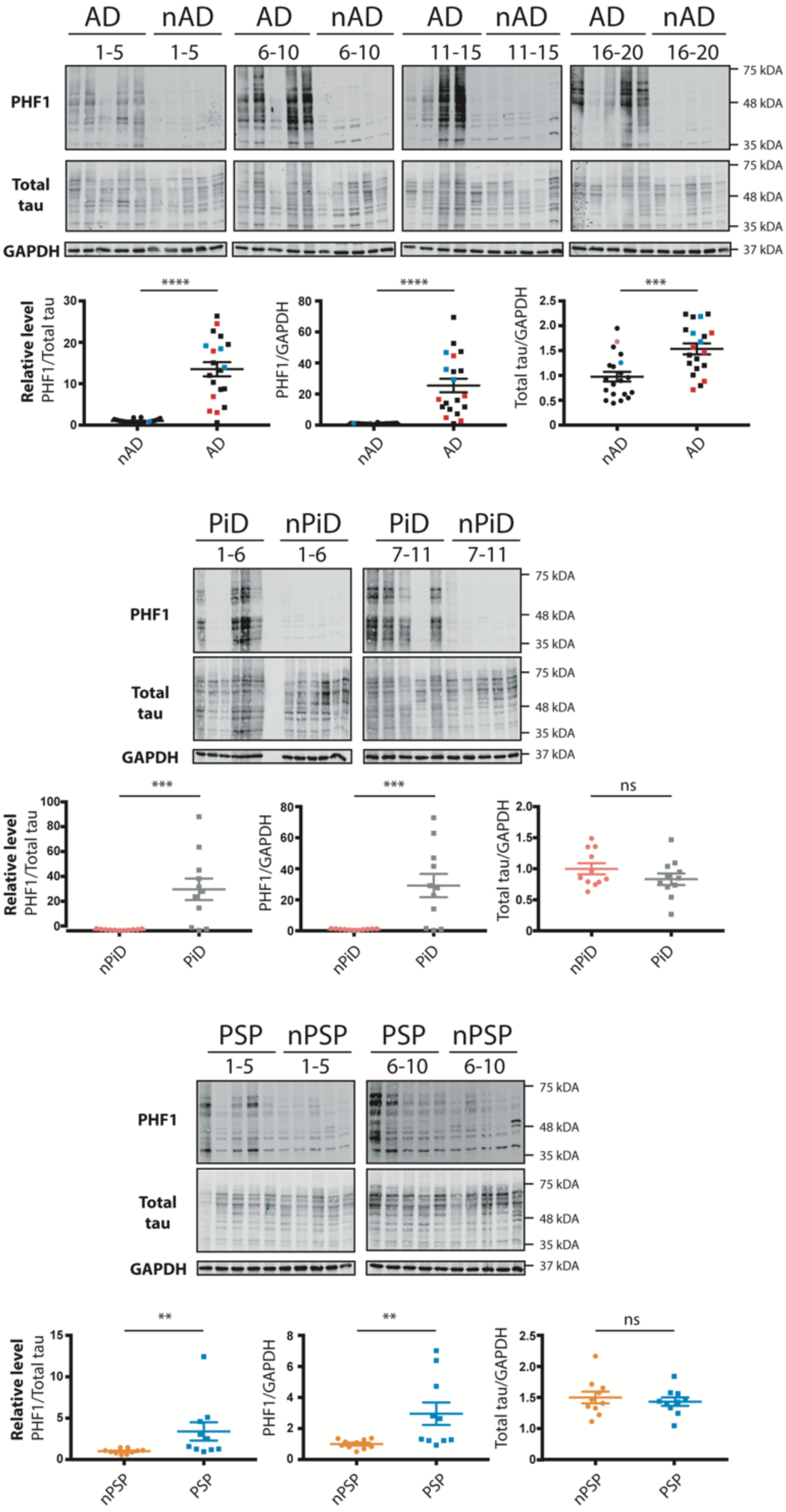
Relative levels of phospho-tau in the brain samples from distinct tauopathy cohorts. Brain homogenates from aged-matched controls and AD, PiD and PSP brains were probed for p-tau, total tau and GAPDH. The PHF1 and total tau immunoreactivity is shown alongside the corresponding GAPDH. Immunoreactivity was expressed as ratios of PHF1/total tau, PHF1/GAPDH and total tau/GAPDH. The normalised data were analysed using one-tailed unpaired t-test (AD and PiD) or Mann-Whitney test (PSP). Total tau was normalised to GAPDH. Data were analysed using two-tailed unpaired t-test (AD and PiD) or Mann-Whitney test (PSP). Error bars are SEM. ****p≤0.0001; ***p≤0.001; **p≤0.01, ns-not significant. Each lane corresponds to one individual case.

### p-eIF2α level in tauopathy

The PERK arm of the UPR has emerged as a candidate stress response in AD and other tauopathies [14,28]. Here we used the phosphorylation of eIF2α that occurs upon PERK activation as a surrogate for the activation of this branch. Importantly, in the preclinical studies an increase in phosphorylation of eIF2α has been discussed as mediating UPR in face of no XBP-1 signalling. The antibody capable of detecting a tunicamycin-induced eIF2α phosphorylation in human cells by Western blot (Supplement S1B) was used to probe whether PERK arm of UPR was activated in humans.

A single or double bands of the correct size were detected for all the samples investigated (Figure 3A). p-eIF2α level was not different between tauopathies and their respective controls (Figure 3B). The absence of increased p-eIF2α suggests that PERK arm of UPR is not activated in the investigated tauopathies.

**Figure 3.**
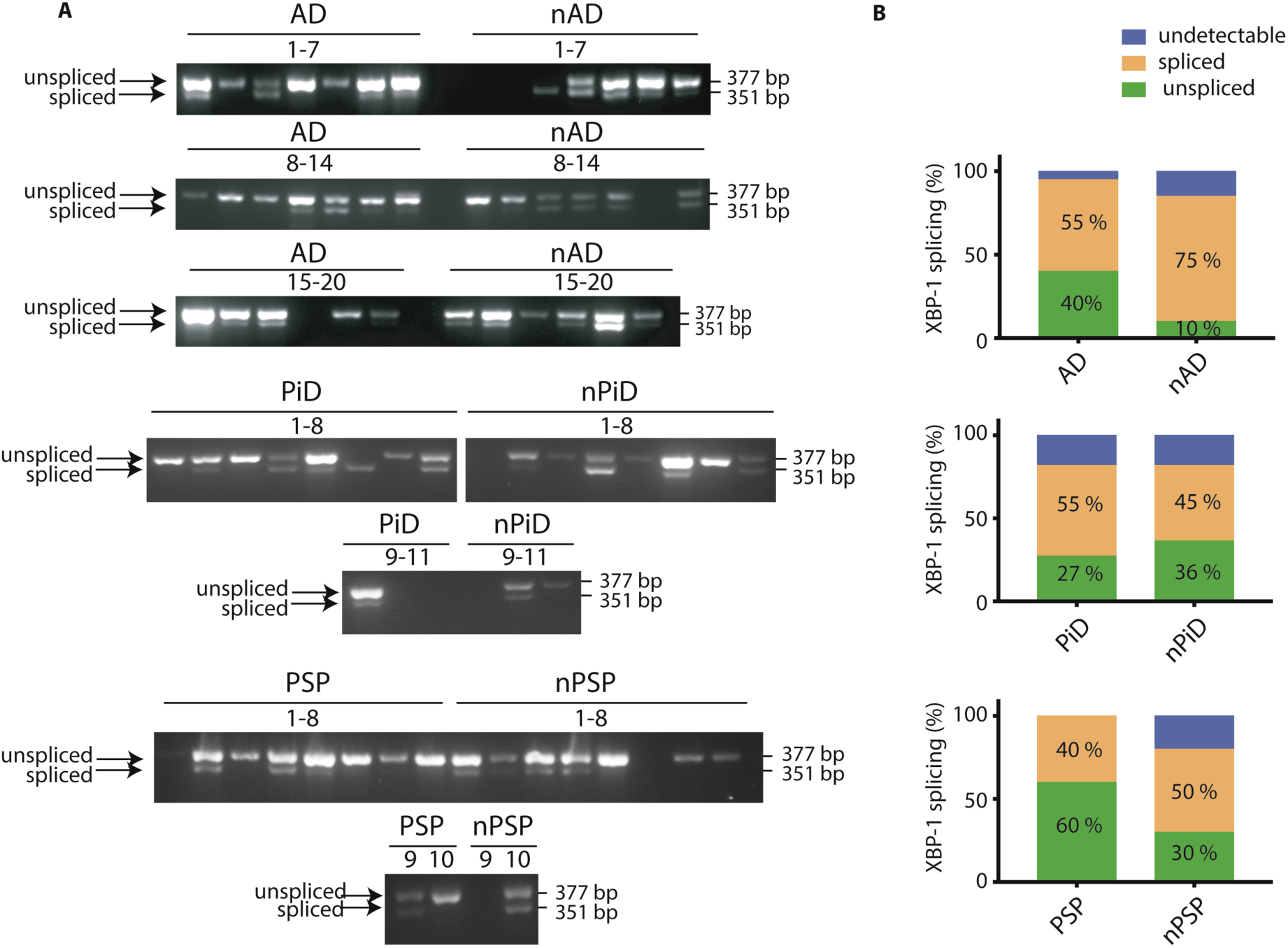
XBP-1 splicing in tauopathies and aged matched control brains. cDNA from brain samples from AD, PiD and PSP together with respective non-demented controls were subjected to PCR with primers designed to amplify the unspliced and spliced variants of XBP-1. The products were resolved on a 2.5% agarose gel (A). The outcome of PCR was classified as undetectable, spliced or unspliced and the distinct PCR outcomes are expressed as percentage of total samples analysed for AD, PiD and PSP samples (B). Each lane corresponds to one individual case.

### Expression levels of the common sensor of the UPR-BiP

BiP is commonly described as a sensor of the UPR and the level of BiP expression is known to be increased upon UPR activation. In contrast to the selective markers described above, p-eIF2α and XBP-1, BiP is implicated in all three branches of the UPR. The detection of BiP expression was performed using previously tested reagents (Supplement S1C and S1D). mRNA expression of BiP did not differ between the disease cases and respective controls (Figure 4A). Similarly, the expression of BiP protein was not changed in the disease vs. control cases (Figure 4B and 4C). The lack of increase of either BiP mRNA or protein suggests that UPR is not activated in the tauopathies.

**Figure 4.**
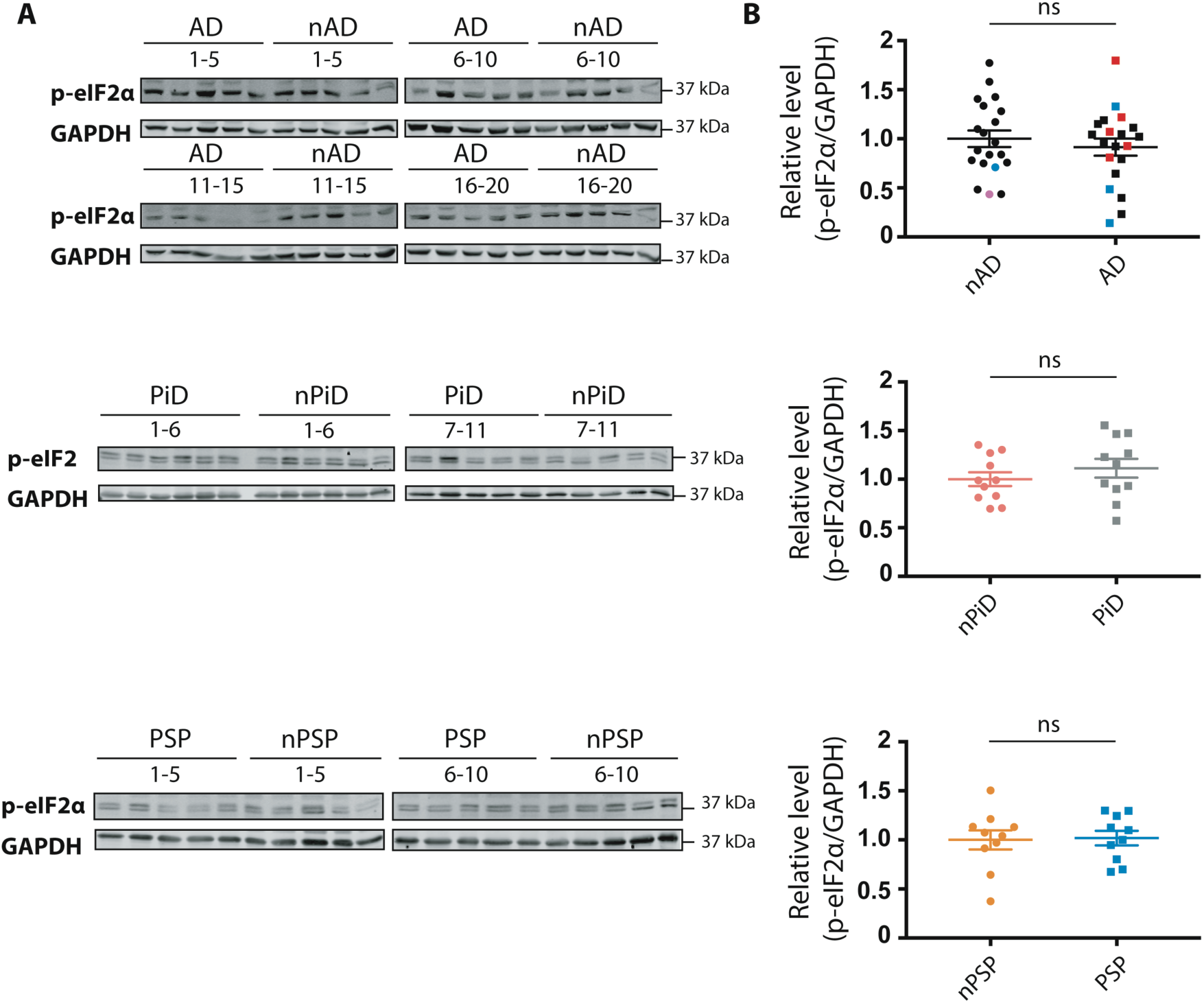
Detection of eIF2α phosphorylation in tauopathies and aged-matched controls. p-eIF2α and GAPDH immunoreactivity for AD, PiD and PSP together with relative age-matched non-demented controls (A). Quantification of p-eIF2α immunoreactivity from individual samples was normalised to the immunoreactivity of a loading control, GAPDH (B). Data were analysed using two-tailed unpaired t-test revealing no significant differences (p= 0.4892, p=0.3604, p=0.8813 for AD, PiD and PSP, respectively). Error bars are SEM. Ns-not significant. Each lane corresponds to one individual case.

### CHOP expression in tauopathies

Finally, CHOP, a sensitive marker that can be induced by all of the three branches of the UPR was investigated. Similar to BiP, mRNA and protein level was determined using previously tested reagents that detected an induced UPR in tunicamycin-treated human cells (Supplement S1E and S1F). The antibody raised against CHOP protein showed the clear induction of a protein of expected size in the tunicamycin-treated cells. However, we noted a confounding co-migration of non-specific bands next to the protein of interest that were enhanced in the untreated cells. To quantify the relative CHOP expression in the brain samples, the immunoreactivity that migrates at the same size of the induced protein in HEK293 cells was used.

No difference in the CHOP mRNA and protein expression was observed between disease and control cases (Figure 5). These findings suggest that the downstream target of all the UPR branches is not induced further supporting the notion that UPR is not activated in the investigated tauopathies.

**Figure 5.**
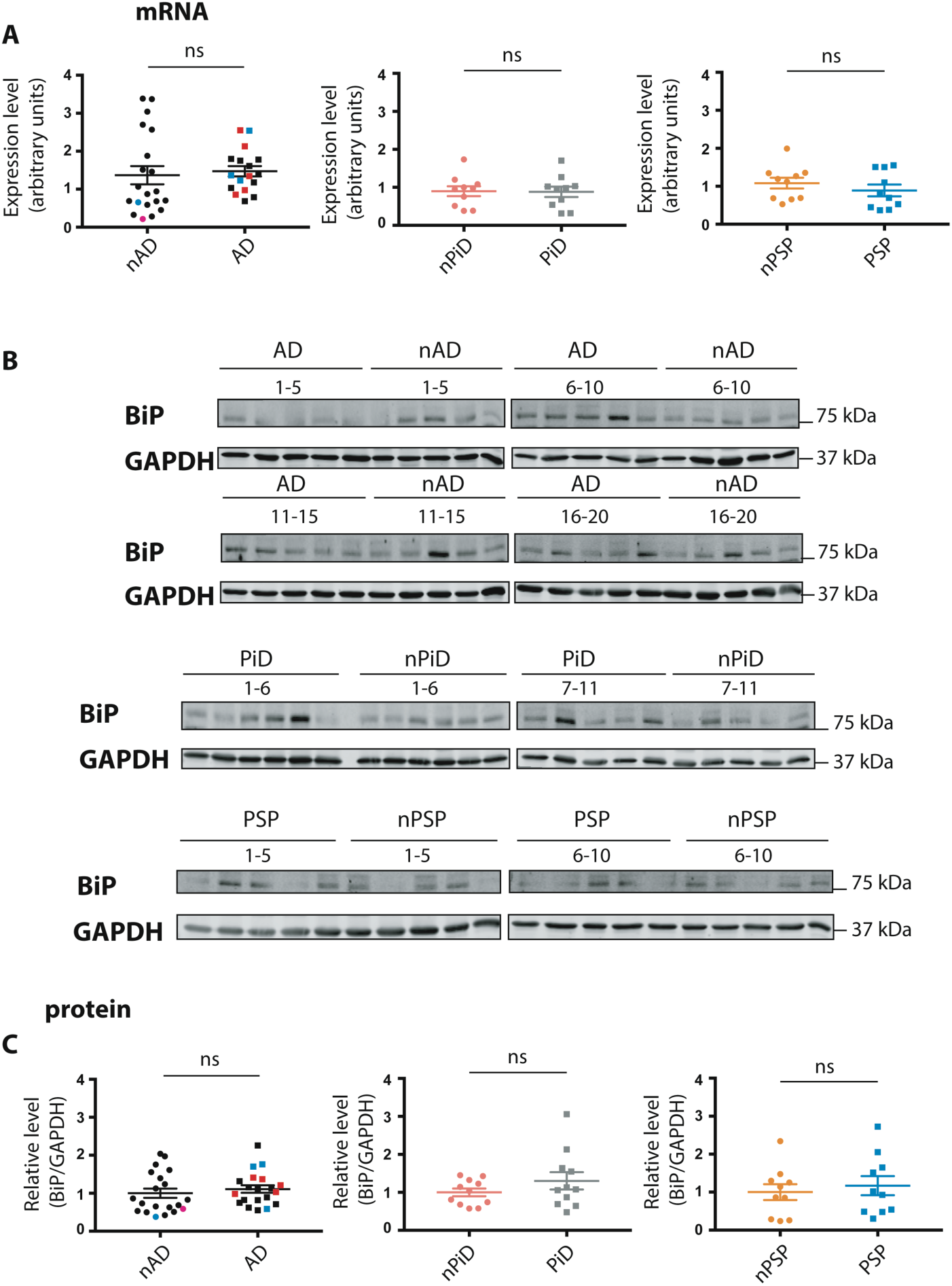
BiP mRNA and protein expression in tauopathies and aged-matched non-demented controls. The normalised expression level of the BiP transcript in AD, PiD and PSP together with respective distinct cohorts of age-matched non-demented controls (A). Data were analysed using Mann-Whitney test (AD) or two-tailed unpaired t-test (PiD and PSP) revealing no significant differences (p= 0.2565, p=0.9356, p=0.3691 for AD, PiD and PSP, respectively). BiP and GAPDH immunoreactivity for AD, PiD and PSP together with relative age-matched non-demented controls (B). Quantification of BiP immunoreactivity from individual samples was normalised to the immunoreactivity of a loading control, GAPDH (C). Data were analysed using Mann-Whitney test (AD) or two-tailed unpaired t-test (PiD and PSP) revealing no significant differences (p= 0.2888, p=0.2402, p=0.6099 for AD, PiD disease and PSP, respectively). Error bars are SEM. Ns-not significant. Each lane corresponds to one individual case.

**Figure 6.**
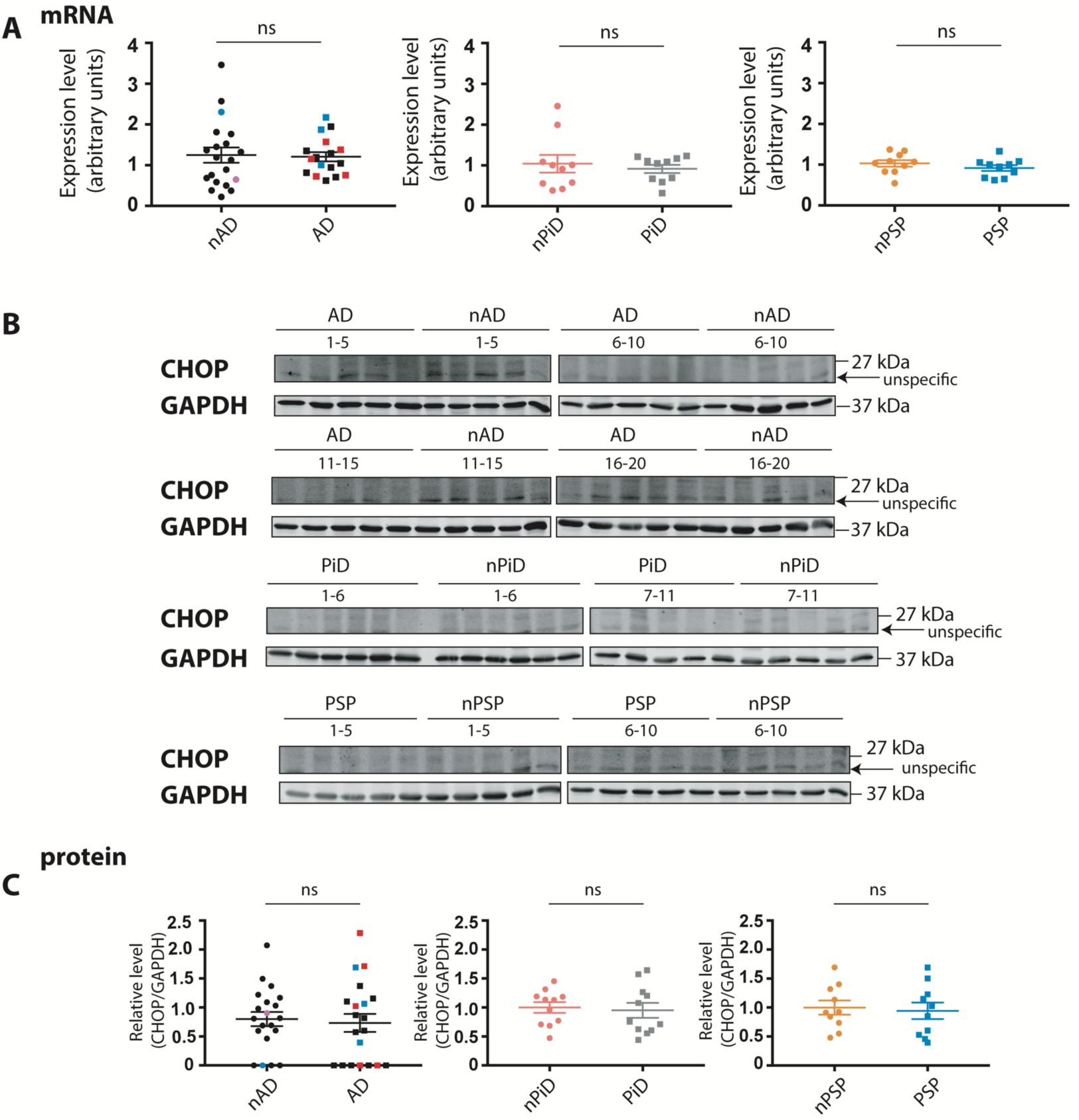
CHOP mRNA and protein expression in tauopathies and aged-matched non-demented controls. The normalised expression level of the CHOP transcript in AD, PiD and PSP together with respective distinct cohorts of age-matched non-demented controls (A). Data were analysed using two-tailed unpaired t-test (AD and PSP) or Mann-Whitney test (PiD) revealing no significant differences (p= 0.8633, p=0.4813, p=0.3043 for AD, PiD and PSP, respectively). CHOP and GAPDH immunoreactivity for AD, PiD and PSP together with relative age-matched non-demented controls (B). Quantification of CHOP immunoreactivity from individual samples was normalised to the immunoreactivity of a loading control, GAPDH (C). Data were analysed using Mann-Whitney test (AD) or two-tailed unpaired t-test (PiD and PSP) revealing no significant differences (p= 0.6749, p=0.7539, p=0.7649 for AD, PiD and PSP, respectively). Error bars are SEM. Ns-not significant. Each lane corresponds to one individual case.

### Correlations of p-tau with the UPR markers

We explored whether the expression of p-tau was associated with the UPR markers within the different groups (disease and controls) of the different tauopathies (AD, PiD and PSP). No significant association was detected between the p-tau markers AT8 and PHF1 with the UPR markers p-elF2a, BIP and CHOP proteins in any of the control or disease groups (Tables 5-7). This again supports the statement that human tauopathies are not associated with the activated UPR.

**Table 5.** Correlations for nAD and AD samples. r_p_- Pearson’s correlation, r_s_- Spearman’s correlation. P values less than 0.01 were considered statistically significant.

| | | PHF1 | AT8 | p-eIF2α | BiP protein | CHOP protein |
| --- | --- | --- | --- | --- | --- | --- |
| PHF1 | nAD | | $r_s=0.237$ | $r_p=-0.284$ | $r_s=0.152$ | $r_p=-0.190$ |
| | AD | | $r_s=0.400$ | $r_p=-0.022$ | $r_p=0.043$ | $r_s=0.459$ |
| AT8 | nAD | $r_s=0.237$ | | $r_s=-0.155$ | $r_s=0.144$ | $r_s=-0.206$ |
| | AD | $r_s=0.400$ | | $r_s=0.059$ | $r_s=0.123$ | $r_s=0.482$ |

**Table 6.** Correlations for nPiD and PiD samples. r_p_- Pearson’s correlation, r_s_- Spearman’s correlation. P values less than 0.01 were considered statistically significant.

| | | PHF1 | AT8 | p-eIF2α | BiP protein | CHOP protein |
| --- | --- | --- | --- | --- | --- | --- |
| PHF1 | nPiD | | $r_s=0.000$ | $r_p=0.299$ | $r_p=-0.228$ | $r_p=-0.037$ |
| | PiD | | $r_s=0.327$ | $r_p=-0.373$ | $r_p=0.685$ | $r_p=0.396$ |
| AT8 | nPiD | $r_s=0.000$ | | $r_s=-0.518$ | $r_s=-0.445$ | $r_s=-0.036$ |
| | PiD | $r_s=0.327$ | | $r_s=-0.464$ | $r_s=-0.136$ | $r_s=-0.236$ |

**Table 7.** Correlations for nPSP and PSP samples. r_p_- Pearson’s correlation, r_s_- Spearman’s correlation. P values less than 0.01 were considered statistically significant.

| | | PHF1 | AT8 | p-eIF2α | BiP protein | CHOP protein |
| --- | --- | --- | --- | --- | --- | --- |
| PHF1 | nPSP | | $r_s=-0.300$ | $r_s=-0.079$ | $r_s=0.115$ | $r_s=0.006$ |
| | PSP | | $r_s=0.745$ | $r_p=-0.327$ | $r_p=-0.399$ | $r_p=0.047$ |
| AT8 | nPSP | $r_s=-0.300$ | | $r_s=0.000$ | $r_s=-0.583$ | $r_s=-0.467$ |
| | PSP | $r_s=0.745$ | | $r_s=-0.212$ | $r_s=-0.200$ | $r_s=0.176$ |

## Discussion

The potential role of the unfolded protein response in neurodegeneration has been highlighted in the experimental models and human samples from neurodegenerative diseases which present disturbed proteostasis. This view has been extended to AD and other tauopathies [14,30,31]. However, the UPR role in neurodegeneration remains puzzling due to the fact that many of the accumulating proteins, including tau are not resident to the ER and do not deposit therein [11]. The UPR involvement in tau-related pathology was previously questioned and investigated and of note, no indication of UPR activation was observed in the transgenic mouse model of tauopathy. No changes in the XBP-1 splicing or the level of p-eIF2α, BiP or CHOP was observed between tau^P301S^-expressing mice and wild type animals [18]. In our recent study, we confirmed the absence of UPR activation in the rTg4510 mouse model expressing tau^P301L^ and we further expanded the UPR investigation to an *in vitro* model of tauopathy [19].

Here, we investigated the activation of UPR in the human tauopathies. We measured the expression level of several UPR markers involved in all three branches of the response: XBP-1 splicing, phosphorylation of eIF2α and the expression of BiP and CHOP. Using reliable reagents, we found no changes in these markers between disease and control cases, nor did we find a correlation with p-tau load. Our findings do not indicate UPR involvement in three distinct diseases-AD, PiD and PSP-with cytosolic tau deposition.

Recently, several studies have highlighted the importance of the ER response in neurodegeneration, mainly focusing on immunohistochemical investigation of UPR branches. In AD, p-PERK immunoreactivity was observed in the hippocampus and occasionally in the temporal cortex whereas no staining was reported in the control cases [29]. In the later study, neuropathological criteria staged the increased p-PERK in AD cases to an early stage of associated tau pathology [14]. Other work in PiD and PSP found that p-PERK and p-IRE1 immunohistochemical staining was highlighted in p-tau positive neurons relative to controls. Similar to the observations from AD brain, the presence of UPR markers was associated with early tau pathology [15]. However, as we note in our study the ability to discern disease selective responses is challenging and it has been found that p-PERK is activated in over 70% of control cases. Interestingly in such studies the increase of UPR markers in the control cohorts correlated to the age of the individual and PHF1 immunolabelling [31]. The conflicting results between the studies mentioned above and data reported in our study could be explained by differences in the disease stage of the investigated cohorts.

Our study failed to show any indication for an activated UPR using the reagents we could verify had requisite specificity. Interestingly, increased level of PERK and p-PERK that was independent of any change in p-eIF2α was reported in PSP. The observation from Bruch et al. regarding lack of the increased eIF2α phosphorylation is consistent with our findings, although we were unable to resolve specific p-PERK immunoreactivity using available reagents.

XBP-1 splicing, a part of IRE1α arm, is a sensitive indicator of UPR activation. It has been previously shown that in the *in vivo* and *in vitro* models of tauopathy, the splicing event does not occur [16,19]. Interestingly, splicing was detected in humans but our findings seem to imply that this event is independent of the disease status. This activation of UPR in diseased and control cohorts is similar to the findings made in investigation of AD and age matched controls [32]. This reinforces the notion that the splicing event is not a disease-specific feature and could be attributed to other factors.

The UPR activation can be also determined using a sensor of UPR, BiP that is upstream of all three branches. There are conflicting results regarding BiP expression in human tauopathies. BiP protein in the brain lysate from the temporal cortex and hippocampus of AD has been reported to increase [29]. However, this contrasts a study that measured BiP level in the cortical homogenates from control, familial and sporadic AD cases. [33] Interestingly, a distinct study reported a significant decrease in the level of BiP in the temporal cortex of sporadic AD cases when compared to controls. What is more, BiP expression was further reduced in the cases with genetic AD [34]. These conflicting results may be due to the cohort size, with ours (n= 20) being the largest relative to the smaller groups used in the above-mentioned studies and/or a consequence of distinct reagents that were used.

Analysing the expression level of the transcripts and corresponding proteins involved in all three arms of UPR, we found no indication of the response activation in three different tauopathies. Interestingly, we observed a high heterogeneity of tau phosphorylation level between the brain samples from patients affected with tauopathies. This high variability is in contrast to the situation observed in transgenic animals in which transgene expression is at the similar level. As the accumulation of p-tau could be a correlate of the phase of the disease, the high variability in p-tau level allowed us to determine whether there is any association between the level of UPR markers and disease stage. We could not find any correlation between the level of p-tau and the UPR markers suggesting that this response is not dependent on tau accumulation.

## Conclusion

In this study, we observe that the accumulation of p-tau did not drive the activation of UPR. Thus, our results call into question the appropriateness of the idea of targeting this response as a possible treatment for tau-related neurodegeneration that has been previously suggested [16,17].

## Acknowledgements

Brain tissue was provided by the London Neurodegenerative Diseases Brain Bank (LNDBB), the South West Dementia Brain Bank (SWDBB) and the Oxford Brain Bank (OBB). All Brain banks are supported by the Medical Research Council (MRC)-UK and the Brains for Dementia Research programme, jointly funded by Alzheimer’s Research UK and Alzheimer’s Society. SWDBB is also supported by BRACE (Bristol Research into Alzheimer’s and Care of the Elderly) and OBB by the NIHR Oxford Biomedical Research Centre.

## Authors contributions

APP, DB, VOC and KD designed the study. APP, IJH performed experiments. APP, IJH and DB analysed the data. APP wrote the manuscript. All authors reviewed and edited the manuscript. VOC and KD acquired funding for the study.

---

**Figure S1.**
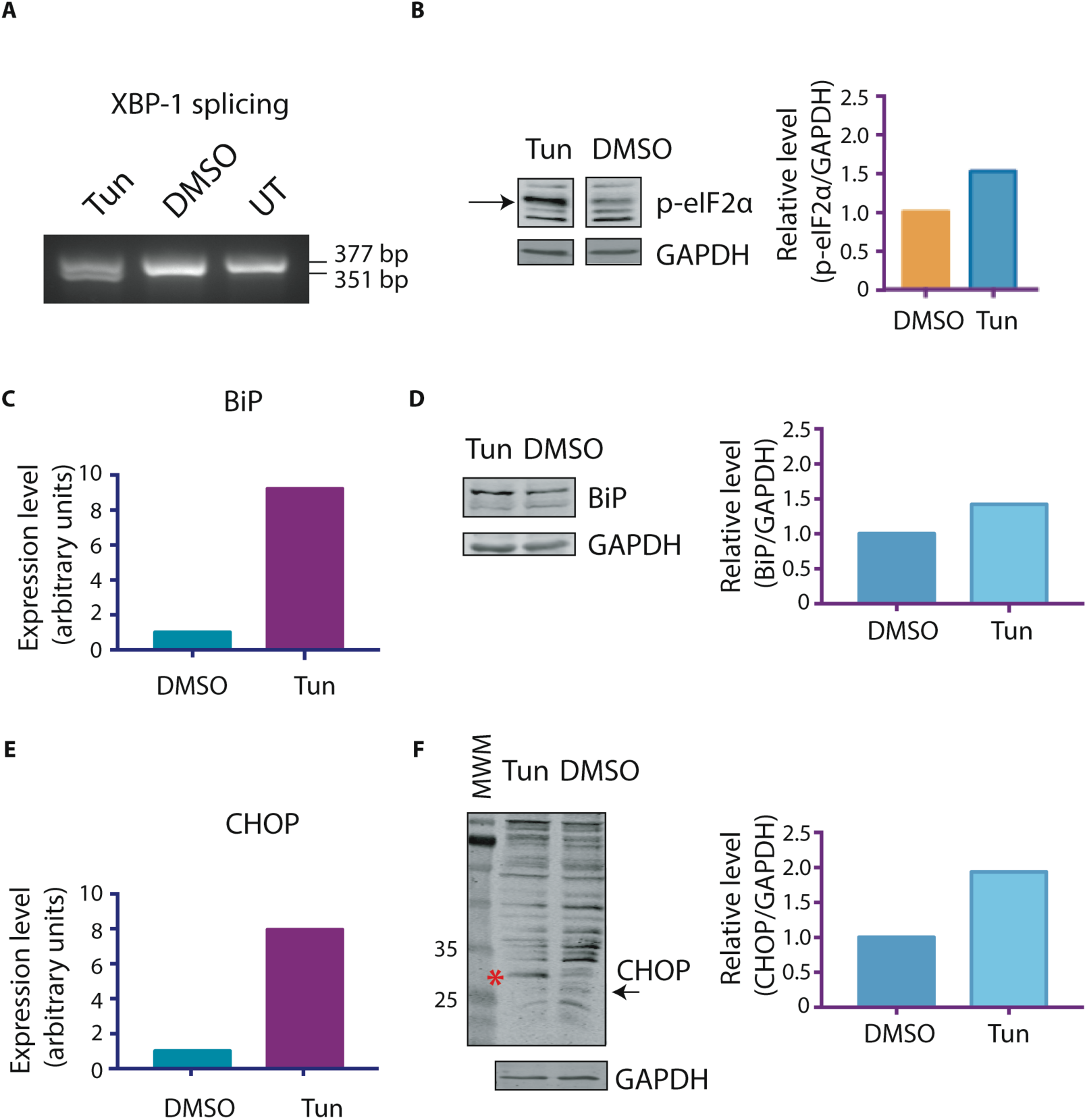
Benchmarking reagents for their ability to resolve UPR in human samples. mRNA and protein were extracted from tunicamycin (Tun, 2 µg/ml)-treated or DMSO (0.1%)-treated cells for 3 (for p-eIF2α detection) or 6 hours. n=1. cDNA from HEK cells showed selective induction of XBP-1 splicing following tunicamycin treatment (A). Modest induction of eIF2α phosphorylation in tunicamycin-treated samples. The samples were reprobed for GAPDH immunoreactivity (B). BiP mRNA and protein expression (normalised to GAPDH) are selectively induced in HEKs treated with tunicamycin (C, D respectively). CHOP mRNA and protein expression (normalised to GAPDH) are selectively induced in HEKs treated with tunicamycin (E, F respectively). Non-specific bands can be observed after the incubation with CHOP antibody that can be seen on the full blot presented. However, tunicamycin treatment leads to the induction of a protein of expected size (indicated with red asterisk) and this band was used for CHOP protein quantification. The arrow indicates the unspecific band that is also highlighted in the figure 5 with an arrow. Molecular weight marker (MWM) highlighting 35 and 25 kDA bands is shown as a reference.

**Figure S2.**
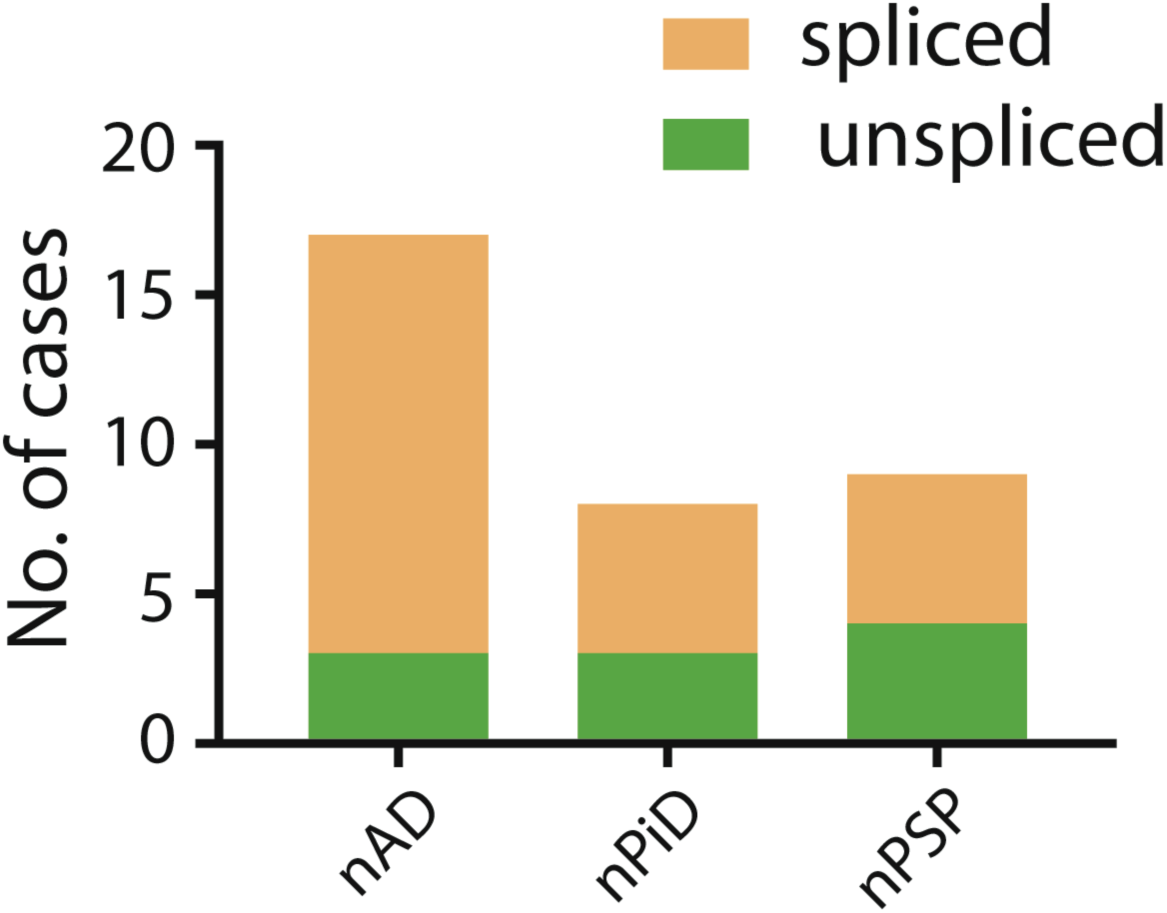
XBP-1 splicing occurs independently of tauopathy. Representation of number of cases that exhibited XBP-1 splicing/lack of splicing within the non-demented cohorts studied. There are no significant statistical differences between the cohorts in terms of XBP-1 splicing (Fisher’s exact test).

